# Apical contacts stemming from incomplete delamination guide progenitor cell allocation through a dragging mechanism

**DOI:** 10.1101/2021.02.24.432660

**Authors:** Eduardo Pulgar, Cornelia Schwayer, Néstor Guerrero, Loreto López, Susana Márquez, Steffen Härtel, Rodrigo Soto, Carl-Philipp Heisenberg, Miguel L. Concha

## Abstract

The developmental strategies used by progenitor cells to endure a safe journey from their induction place towards the site of terminal differentiation are still poorly understood. Here we uncovered a progenitor cell allocation mechanism that stems from an incomplete process of epithelial delamination that allows progenitors to coordinate their movement with adjacent extra-embryonic tissues. Progenitors of the zebrafish laterality organ originate from the surface epithelial enveloping layer by an apical constriction process of cell delamination. During this process, progenitors retain long-term apical contacts that enable the epithelial layer to pull a subset of progenitors along their way towards the vegetal pole. The remaining delaminated progenitors follow apically-attached progenitors’ movement by a co-attraction mechanism, avoiding sequestration by the adjacent endoderm, ensuring their fate and collective allocation at the differentiation site. Thus, we reveal that incomplete delamination serves as a cellular platform for coordinated tissue movements during development.

**Impact Statement:** Incomplete delamination serves as a cellular platform for coordinated tissue movements during development, guiding newly formed progenitor cell groups to the differentiation site.

## Introduction

During embryo development, naïve cell lineages undergo concurrent processes of fate specification and morphogenesis as critical steps towards the generation of differentiated tissues and organs. These early progenitor cells often ought to travel long distances from their induction site to the site of terminal differentiation, making them vulnerable to environmental signals and the movement of neighbouring tissues that may impede a correct path or change their potential, and consequently reduce the pool of progenitors available for subsequent stages of differentiation. The success of this journey is especially important when the number of progenitors of a tissue or organ is limited. In these cases, small reductions in the number of progenitors can lead to developmental abnormalities that result in a dysfunctional organ (Moreno-Ayala, et al., 2020). Various embryonic tissues and organs originate from small sets of progenitor cells, including the primordial germ cells that give rise to gametes in the gonads of vertebrates and invertebrates, the primordia of the posterior lateral line that give rise to neuromasts along the trunk and tail of fish and amphibians, and the progenitors of the laterality organ that participate in left-right pattern formation in several vertebrates (reviewed in Dalle Nogare and Chitnis, 2017; Reig, et al., 2014; Matsui and Bessho, 2012; Richardson and Lehmann, 2010). Despite the importance of the developmental paths followed by these small groups of progenitor cells and their impact on the physiology of the organism we still know little about the array of developmental strategies progenitors cells deploy *in vivo* to overcome the challenges imposed by the environment while travelling to the site of terminal differentiation. Here we examine this question during the early stages of morphogenesis of the embryonic laterality organ, the first organ to be formed during vertebrate development, using zebrafish as a model organism.

The laterality organ or left-right organiser of zebrafish is a transient embryonic structure of epithelial nature known as the Kupffer’s vesicle that contains motile cilia required for the determination of the left-right axis (Essner, et al., 2005; Kramer-Zucker, et al., 2005; Cooper and D’Amico, 1996). This organ-like epithelial structure originates from a small group of 20-30 progenitors, known as dorsal forerunner cells (DFCs), which arise at the dorsal margin of the late blastula by a process of cell ingression that converts the superficial epithelial cells of the extra-embryonic enveloping layer (EVL) into deep mesenchymal-type DFCs (Oteiza, et al., 2008). After ingression, DFCs move as a cellular collective from their place of origin at the equator of the embryo towards the vegetal pole, reaching the terminal location for organ differentiation at the posterior tip of the notochord (Figure 1A; Supplementary Video 1) (Oteiza, et al., 2008; Cooper and D’Amico, 1996). On their travel to the site of differentiation, DFCs are located ahead of the margin of the deep cell layer (DCL) where specification signals and massive internalisation movements transform the marginal epiblast into the mesendoderm (Figure 1B) (Pinheiro and Heisenberg, 2020). Despite the proximity to the DCL margin and the fact that DFCs share critical determinants with the mesendoderm (Warga and Kane, 2018; Alexander and Stainier, 1999), the movement and fate of DFCs do not seem to be affected by the specification signals nor by the *in mass* internalisation movements of the mesendoderm, remaining separated from this cellular domain during their vegetal movement. On the other hand, DFCs follow the same vegetal ward direction of movement as the overlying EVL during epiboly (Bruce and Heisenberg, 2020) and appear to be physically connected with this extra-embryonic epithelial tissue as revealed by the presence of puncta enriched in the TJ-associated zonula occludens 1 protein (ZO-1) at the DFC-EVL interface (Ablooglu, et al., 2010; Oteiza, et al., 2008). This observation raises the question of whether the presumed DFC-EVL connections play a role in the movement of DFCs towards the organ differentiation site, a hypothesis that has not yet been tested experimentally.

**Figure 1.**
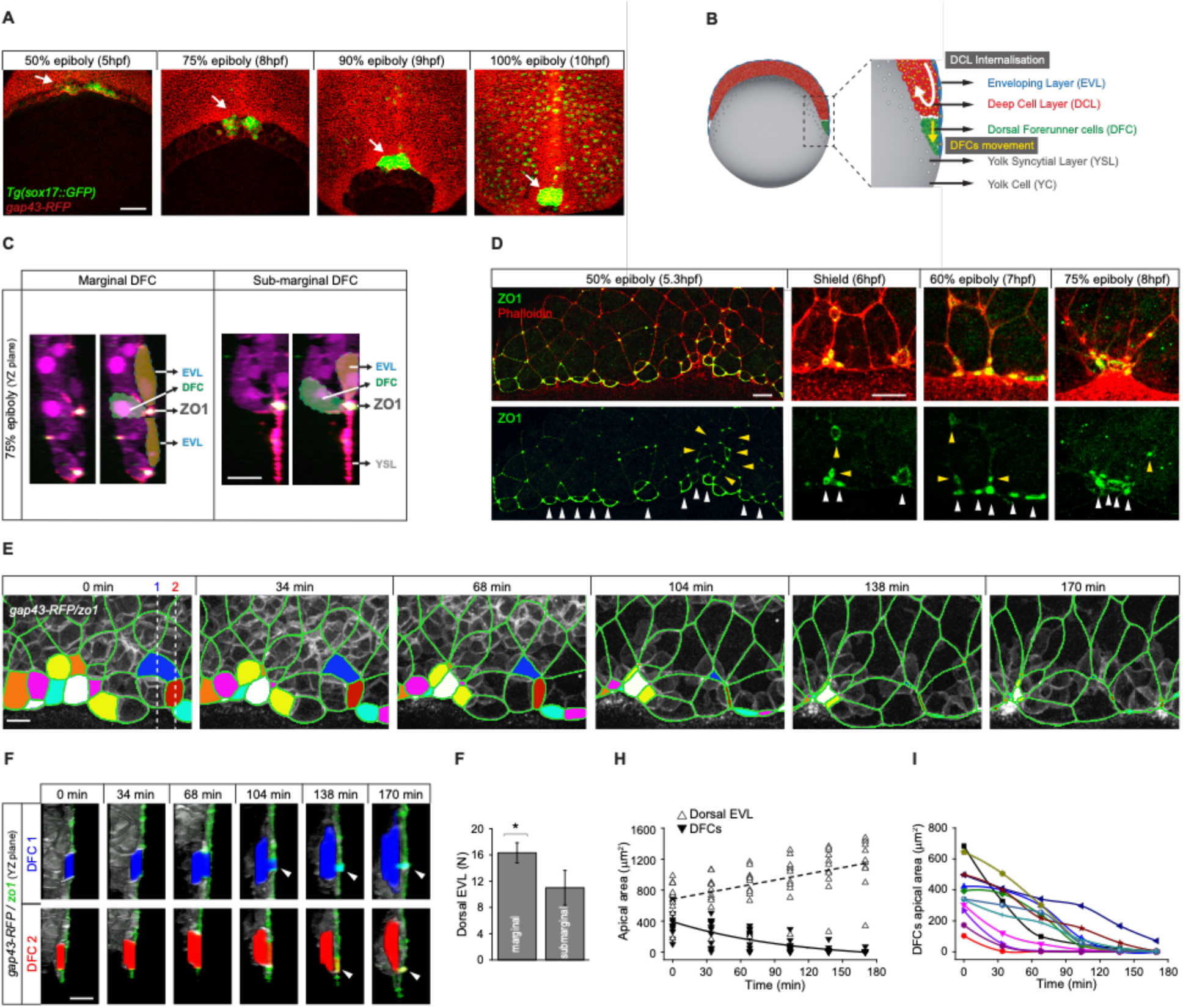
DFCs delaminate by apical constriction and retain apical attachments with the EVL and YSL. (A) Dorsal views of confocal z-stack maximum projections showing the collective vegetal motion of DFCs between shield stage and 100% epiboly in a representative Tg(*sox17::GFP*) embryo injected with *gap43-RFP* mRNA. DFCs are in green (arrows) while the plasma membrane of all cells is in red. Note that the *sox17::GFP* transgene also labels the scattered population of endodermal cells at advances stages of epiboly (extracted from Supplementary Video 1). Scale bar, 100 µm. (B) Schematic diagram of a cross section along the sagittal plane of the zebrafish embryo at 60% of epiboly. DFCs move to the vegetal pole ahead of the DCL margin, where mesendodermal progenitors internalise. (C) Confocal microscopy zy-plane in 75% epiboly embryos stained with phalloidin and ZO-1 (merge channels), showing marginal (left) and submarginal (right) ingressing DFCs connected with the EVL and YSL by focal apical attachments enriched in ZO-1 and F-actin. Scale bars, 20 µm. (D) Phalloidin and ZO-1 immunostaining (merge on top and ZO-1 on bottom) of the dorsal margin of wild type embryos between 50% and 75% epiboly. Images correspond to surface confocal sections showing the apical domains of delaminating DFCs in contact with the EVL and YSL (arrowheads). Scale bar, 20 µm. (E) Time series of dorsal views of confocal z-stack maximum projections of a representative embryo injected with *zo1-GFP* and *gap43-RFP* between 50% and 80% epiboly, showing EVL cell junctions (green outlines) and the apical domains of EVL cells as they delaminate to become DFCs (coloured areas) (extracted from Supplementary Video 2). Scale bar, 20 µm. (F) Time series of confocal z-sections showing two DFCs taken from panel E in lateral views (blue and red cells) as they move below the plane of the EVL epithelium during the process of delamination. Note that delaminating DFCs retain a focal apical attachment with the EVL (arrowhead, top) and YSL (arrowhead, bottom). Scale bars, 20 µm. (G) Quantification of the number of dorsal EVL cells undergoing delamination to become DFCs at both marginal and submarginal positions, expressed as means ± s.d. * (*p*<0.05). (H) Temporal changes in apical area of DFCs (black triangles) and neighbouring dorsal EVL cells (white triangles) in a representative embryo during the process of delamination. Continuous and dashed lines indicate the mean values of apical area of DFCs and dorsal EVL cells, respectively. (I) Temporal changes in apical area of individual DFCs in a representative embryo during the process of delamination. Each curve corresponds to a single cell. Animal is to the top in all panels.

Here we addressed this hypothesis by combining in vivo imaging and biomechanical manipulation in zebrafish embryos. We found that DFCs arise from the EVL by an actomyosin-mediated apical constriction process of cell delamination. During this process, DFCs retain long-term physical connections with the EVL and yolk syncytial layer (YSL) through tight junction-enriched apical attachments, at the time both extra-embryonic tissues spread to the vegetal pole in the movement of epiboly. We demonstrated that extra-embryonic tissue spreading is transmitted to DFCs through the apical attachments to drag their vegetal movements. As DFCs detach from the extra-embryonic cellular domains after completing delamination or undergoing division, polarised protrusions and E-cadherin mediated adhesion integrate detached cells to the vegetal movement of attached DFCs, avoiding sequestration by the endoderm and ensuring the vegetal motion of all progenitors as a group. Thus, we unveil a drag-mediated guidance mechanism of progenitor cell morphogenesis that relies on the concerted activities of the process of epithelial delamination underlying progenitor cell specification and the directed spreading of adjacent extra-embryonic tissues.

## Results

### DFCs ingress by cell delamination through an apical constriction process that provides long-term apical attachments with extra-embryonic tissues

The previous observation of puncta enriched in the tight junction adaptor protein ZO-1 at the interface between a subset of DFCs and the overlying EVL during late epiboly stages (Ablooglu, et al., 2010; Oteiza, et al., 2008) suggests that DFCs establish contacts that attach them to the extra-embryonic surface epithelium. To investigate this possibility, we started by studying the spatial distribution of these enriched ZO-1 regions. Double immunolabelling for ZO-1 and phalloidin at 75% epiboly confirmed that ZO-1 puncta marked discrete regions, also enriched in F-actin, where DFCs contacted the EVL at cell-cell junctions near the margin of the epithelium (Figure 1C, left). Also, we observed the punctuated co-accumulation of ZO-1 and F-actin at the epithelial margin where DFCs contacted the YSL (Figure 1C, right). Indeed, the latter population of marginal DFCs was more abundant than submarginal DFCs (Figures 1D and 1G). We then asked how these contacts arise by examining the temporal progression of ZO-1 from early to late epiboly stages throughout DFC formation. Previous work have shown that DFC originate from dorsal EVL cells by a process of ingression (Oteiza, et al., 2008) although the underlying mechanism is currently unknown. We found that the punctuated ZO-1 and F-actin staining observed at late epiboly stages resulted from apical area reduction as DFCs underwent ingression from the EVL. At early epiboly stages, ZO-1 accumulated along the apical junctions of DFC progenitors (Figure 1D). As epiboly progressed, the apical face of these cells gradually reduced in area to leave punctuated apical domains enriched in ZO-1 and F-actin proteins (Figure 1D). Together, these findings reveal that DFCs ingress from the EVL by a mechanism of cell delamination mediated by apical constriction. Remarkably, this process leaves discrete apical attachments with the extra-embryonic tissues.

To investigate the dynamics of apical constriction leading to the formation of apical attachments we performed *in vivo* imaging from late blastula to epiboly stages using a GFP-tagged version of ZO-1. We found that EVL cells fated to become DFCs gradually shrunk their apical area while moving beneath the epithelial sheet (Figures 1E and 1F; Supplementary Video 2), a behaviour that was specific as neighbouring EVL cells not fated to become DFCs exhibited the opposite behaviour, increasing their apical area as epiboly progressed (Figure 1H). Interestingly, the kinetics of apical area reduction varied among DFCs (Figure 1I) and resulted in discrete apical attachments that connected these cells with the YSL and EVL for extended periods (Figure 1F). Collectively, these findings indicate that DFCs retain long-term apical attachments with the extra-embryonic YSL and EVL as a consequence of the apical constriction process of cell delamination that gives rise to the cells.

### DFC apical constriction depends on actomyosin dynamics

Contractile forces generated by an apical actomyosin network have been implicated in driving apical constriction in a variety of developmental processes (Heer and Martin, 2017). We thus assessed if actomyosin dynamics was responsible for DFC apical constriction by performing live imaging using a double fluorescent reporter line for non-muscular (NM) myosin II and F-actin. We found that apical constriction was accompanied by the medioapical accumulation of myosin II and F-actin at the apical face of DFCs (Figure 2A). Quantification of fluorescent intensities revealed that myosin II and F-actin accumulations increased as apical constriction progressed during DFC delamination (Figure 2B). The kinetics of medioapical actomyosin accumulation was asynchronous among DFCs (Figure 2B) reflecting the variation in the kinetics of apical area reduction previously observed within the DFC population (Figure 1I). Importantly, although the extent of medioapical myosin II and F-actin protein accumulation was variable among DFCs (Figure 2B) we found that it scaled with the reduction of DFC apical area (Figures 2C and 2D) suggesting that actomyosin contractility mediates DFC apical constriction. To investigate this possibility, we performed immunostaining for phospho-myosin II light chain, as phosphorylation of the myosin II light chain subunit regulates actomyosin contractility (Heissler and Sellers, 2016). We found that phospho-myosin II light chain was distributed in the apical face of DFCs during apical constriction confirming the contractile nature of the apical actomyosin network (Figures 2E and 2F). To confirm that actomyosin dynamics is required for apical constriction we studied the cases of DFCs showing a transitory increase of apical area while undergoing apical constriction. We observed a sequence of concatenated changes in apical area and myosin accumulation. First, apical constriction was concurrent with the accumulation of myosin at the apical face of DFCs (Figures 2G, 7 min, and 2H, yellow dotted line; Supplementary Video 3). Then, the loss of apical myosin was synchronous with the increase of DFC apical area (Figures 2G, 18 min, and 2H, light blue dotted line; Supplementary Video 3). Finally, the recovery of apical myosin was followed by constriction of DFC apical area (Figures 2G, 24 min, and 2H; Supplementary Video 3). Together, these findings indicate that a contractile medioapical actomyosin network regulates the apical constriction process that mediates DFC delamination.

**Figure 2.**
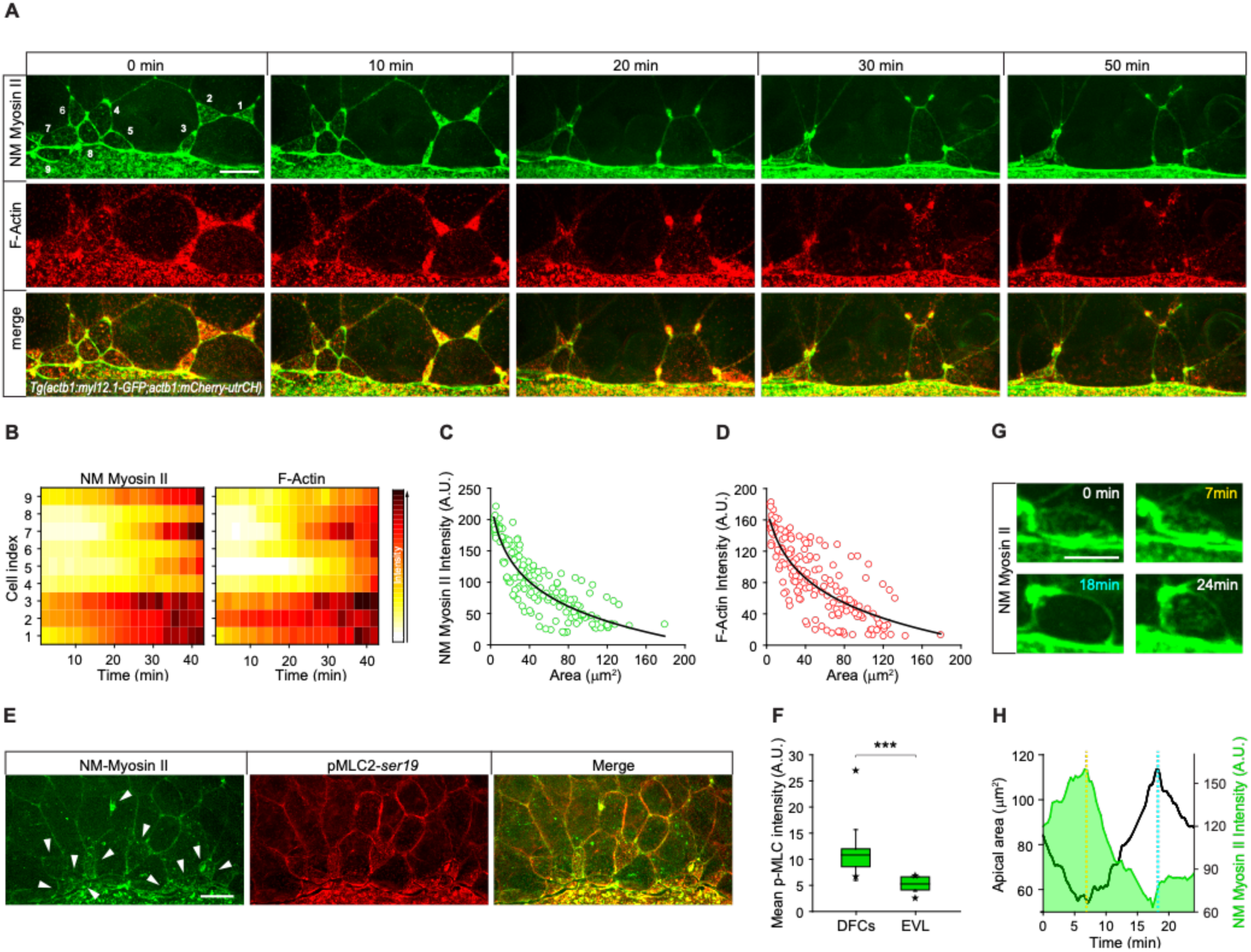
DFC apical constriction is mediated by actomyosin dynamics. (A) Time series of dorsal views of confocal z-stack maximum projections of a representative Tg(*actb1:myl12.1-GFP;lifeactin-RFP*) embryo starting at shield stage, showing the progressive accumulation of NM myosin II (green, top) and F-actin (red, middle) at the apical surface of delaminating DFCs. The merge image of NM myosin and F-actin is shown at the bottom. Scale bar, 20 µm. (B) Heat maps showing the temporal changes in the fluorescence intensity levels of apical NM myosin II (left) and F-actin (right) for individual DFCs in a representative Tg(*actb1:myl12.1-GFP;lifeactin-RFP*) embryo, according to the colour scale shown on the right. Each value corresponds to the average fluorescent intensity for a single cell over a 2.1 min time window. (C and D) Changes in the mean fluorescent intensity levels per pixel of apical NM myosin II (C) and F-actin (D) as a function of the changes in DFC apical area during the process of DFC apical constriction. Dots correspond to values of individual DFCs and lines to the fitted curves, taken from a representative Tg(*actb1:myl12.1-GFP;lifeactin-RFP*) embryo.(E) Dorsal views of the margin of Tg(*actb1:myl12.1-GFP*) embryos at 75% epiboly immunostained to reveal total NM myosin II (green, left), phospho-myosin light chain II (middle) and merge (right). Arrowheads point to the apical surface of delaminating DFCs. Scale bar, 20 µm. (F) Quantification of the fluorescent intensity levels of phospho-myosin light chain II at the apical surface of DFCs and neighbouring dorsal EVL cells. The box depicts the interquartile range from 25% to 75% of the data around the average (vertical line inside the box), the whisker depicts s.d., and stars indicate maximum and minimum values (n=3 embryos). *** (*p* < 0.001). (G) Example of a representative DFC showing spontaneous consecutive apical loss and recovery of myosin II accumulation during the process of apical constriction (extracted from Supplementary Video 3). Scale bar, 10 µm. (H) Quantification of the temporal evolution of apical area and myosin II intensity of the cell shown in G, during the consecutive stages of loss and recovery of apical myosin II. The dashed yellow and light blue lines depict the minimum and maximum values of apical area, which coincide with the maximum and minimum values of apical area during the period of analysis, respectively.

### Cell delamination is concurrent with the vegetal movement of DFCs and extra-embryonic tissues

During epiboly, DFCs move from the embryo equator to the vegetal pole where they differentiate into the laterality organ (Oteiza, et al., 2008). Previous studies have hypothesised that the vegetal movement of these cells relies on polarised protrusive activity (Zhang, et al., 2016; Ablooglu, et al., 2010). However, the presence of apical attachments connecting DFCs with the YSL and EVL opens the possibility that extra-embryonic tissues guide the vegetal movement of DFCs. To discriminate between these two guidance mechanisms, we first examined the organisation of polarised protrusions in DFCs. Using *in vivo* imaging in an actin transgenic reporter line we found that DFCs formed dynamic membrane protrusions (Supplementary Figure 1A), as previously observed (Zhang, et al., 2016; Ablooglu, et al., 2010). However, protrusions were enriched at the rear (lateral and animal) edges and not at the leading (vegetal) edge of the DFC cluster (Figures S1B-S1E) arguing against a primary role of polarised cell protrusions in directing the vegetal movement of DFCs. Therefore, DFC apical attachments could play a key role in guiding the vegetal movement of these cells. To experimentally address this hypothesis, we first determined the extent to which the vegetal movement of DFCs temporally matched the process of delamination using confocal microscopy (Figure 3A). We observed that DFCs travelled over 350 µm towards the vegetal pole from the time in which the first DFCs started the process of apical constriction at around dome stage until the last cells delaminated after detachment from the extra-embryonic tissues after 90% epiboly (Figure 3B, horizontal axes). In agreement with the variable kinetics of apical constriction (Figure 1I) and actomyosin dynamics (Figure 2B), the onset of apical constriction and the time of detachment leading to delamination were also highly variable in the DFC population (Figures 3B, red and blue circles, and 3C). Together, the absence of vegetal ward polarised cell protrusions in DFCs, and the close concurrency of cell delamination with the vegetal movement of DFCs suggest that apical attachments with the YSL and EVL integrate DFCs to the epiboly movements of the extra-embryonic tissues. In support of this idea, the simultaneous tracking of DFCs and EVL cells revealed that the speed, direction and extent of DFC vegetal movement correlated with the vegetal spreading of the EVL (Figures S2A-S2C). Furthermore, DFCs and overlying EVL cells shared the same orientational bias in the axis of cell elongation (Supplementary Figure 2D) indicating a morphogenetic coupling between the two tissues. Collectively, these results suggest that apical attachments derived from the process of cell delamination work as tissue-tissue connectors that couple DFCs with the vegetal spreading of the extra-embryonic YSL and EVL, guiding their vegetal movement during epiboly.

**Figure 3.**
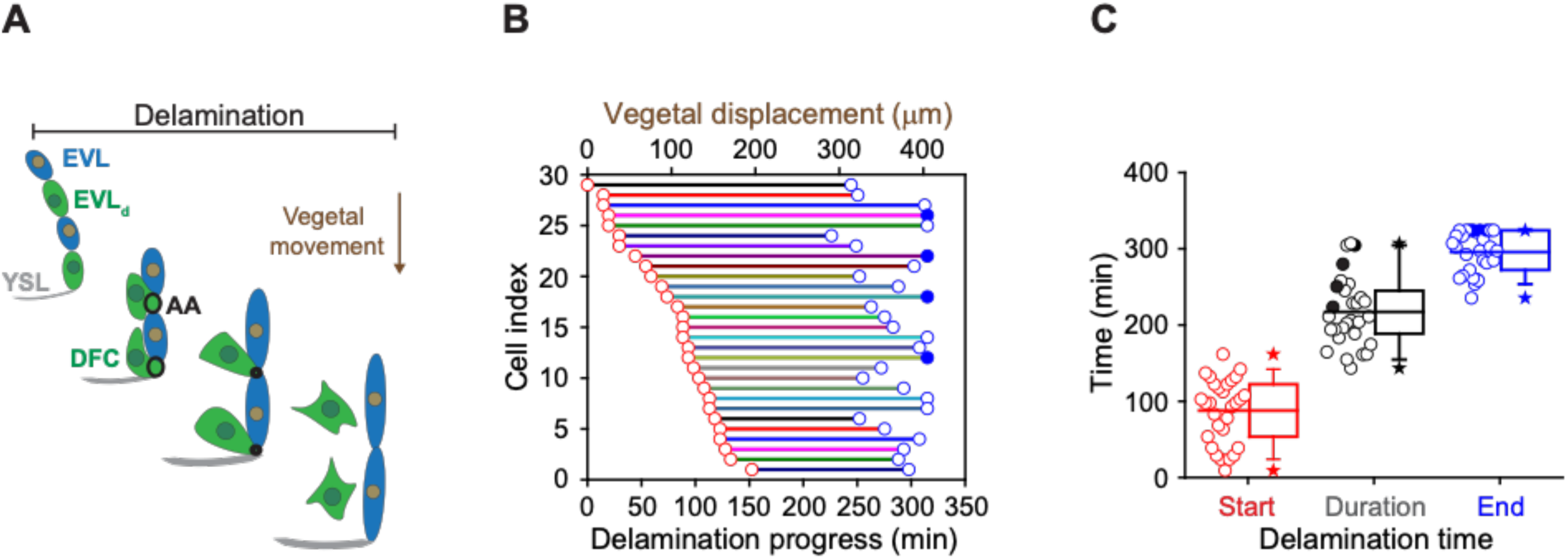
DFC delamination is asynchronous and coexists with the vegetal motion of DFCs. (A) Schematic diagram showing the origin of DFCs from the EVL through cell delamination. Apical attachments (AA) that result from apical constriction connect DFCs with the EVL and YSL during their vegetal movements. When DFCs complete delamination, they loose apical attachments and are released from the EVL. (B) Plot depicting the relationship between the time of the delamination process and the vegetal movements of DFCs for a single representative Tg(*actb1:lifeactin-RFP*) embryo. The start (red circles) and end (blue circles) times of cell delamination, and the total duration of this process (horizontal lines) are shown for individual DFCs (bottom axis) and compared with their vegetal movements (top axis). Filled blue circles indicate a subset of DFCs that still retained apical attachments by the end of the movies. (C) Combined box and distribution plots of the start and end times, and of the total duration of cell delamination for the same representative embryo as in B. Circles correspond to individual values while the box depicts the interquartile range from 25% to 75% of the data around the average (vertical line inside the box), the whisker depicts s.d., and stars indicate maximum and minimum values. Filled blue and black circles indicate the subsets of DFCs that still retained apical attachments by the end of the movies.

### Extra-embryonic tissues pull DFCs through apical attachments to guide their vegetal movement

To directly test if apical attachments between DFCs and the extra-embryonic tissues guide the vegetal movement of DFCs we first disrupted the vegetal spreading of the YSL/EVL. Previous reports have shown that contraction and friction-based flows of a ring-like YSL actomyosin network drive YSL/EVL vegetal spreading during epiboly (Heisenberg and Bellaiche, 2013; Behrndt, et al., 2012; Cheng, et al., 2004). Therefore, we inhibited the activity of the YSL actomyosin ring by decreasing the levels of phosphorylated myosin II through expressing, specifically in the YSL, the N-terminal region of the myosin phosphatase target subunit 1 (N-ter-MYPT1) (Figures S3A and S3B). Embryos expressing N-ter-MYPT1 (MYPT1) showed a speed drop in YSL/EVL vegetal spreading. Likewise, DFCs decreased their vegetal ward speed to the same extent (Figures S3C and S3D). Similarly, local disruption by laser ablation of the YSL actomyosin ring in direct contact with marginal DFCs decreased the speed of EVL spreading and DFC movement near the ablated zone (Figures S3E and S3F). Together, these results demonstrate that the progress of DFC vegetal movement requires the vegetal spreading of the extra-embryonic YSL/EVL.

Next, we conducted two types of experiments to assess if extra-embryonic tissue movement is transmitted to DFCs through their apical attachments. As a first experiment, we disrupted the animal-vegetal tension exerted by EVL spreading on the apical attachments through ablating a cortical EVL junction next to DFC apical attachments (Figure 4A). After ablation, apical attachments recoiled towards the animal pole triggering a fast retraction of DFCs in the same direction (Figures 4B and 4C; Supplementary Video 4). Subsequently, the constricted cortical zone induced by wound repair pulled apical attachments towards the vegetal pole, restoring the vegetal directionality of DFC movement (Figures 4B and 4C; Supplementary Video 4). Finally, we analysed the few cases in which single DFCs developed in isolation far from the central cluster of DFCs. After tracking single isolated cells, we observed that DFCs devoid of apical attachments (hereafter referred to as “detached DFCs”) moved without persistence and directionality, even towards the animal pole in the opposite direction to the vegetal movement of the YSL/EVL (Figures 4D-4G; Supplementary Video 5). In contrast, DFCs connected with the extra-embryonic tissues by apical attachments (hereafter referred to as “attached DFCs”) showed a persistent and directed movement toward the vegetal pole, mimicking the movement of extra-embryonic tissues (Figures 4D-4G; Supplementary Video 5). Collectively, these findings demonstrate that apical attachments transmit the vegetal spreading of extra-embryonic tissues to attached DFCs guiding their vegetal movement during epiboly.

**Figure 4.**
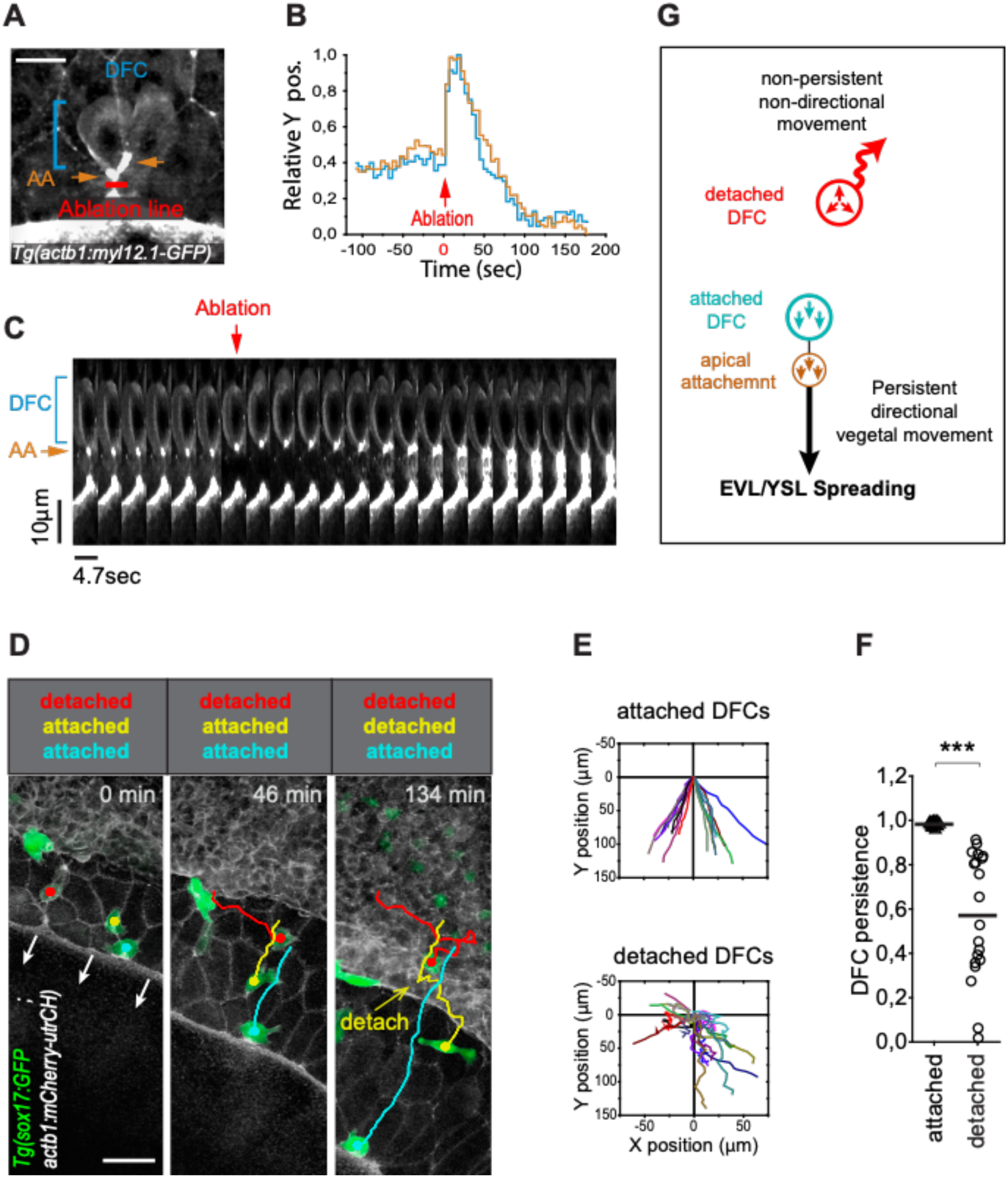
Apical attachments transmit extra-embryonic tissue spreading to guide DFC vegetal movement. (A-C) Laser line ablation of an EVL cortical junction below the apical attachments of two submarginal DFCs in a Tg(*actb1::myl12.1-GFP*) embryo at 70% of epiboly. (A) Dorsal view of a pre-ablation stage showing DFCs (blue bracket), apical attachments (AA, orange arrows) and the ablation line (red). (B) Plot showing the relative position of DFCs (blue) and apical attachments (orange) along the y-axis before and after ablation, with zero corresponding to the time of the laser pulse (red arrow). (C) Kymograph of DFC and apical attachment movements during the laser ablation plotted in B (extracted from Supplementary Video 4). Scale bars, 20 µm. (D-F) Tracking of isolated DFCs obtained from dorsal views of confocal z-stack maximum projections of living 60% epiboly Tg(*sox17::GFP;actb1::mCherry-utrCH*) embryos (D, extracted from Supplementary Video 5). Plots show the differences in movement directionality (E) and persistence (F) between attached and detached DFCs (n=22 attached DFCs from 11 embryos, 20 detached DFCs from 8 embryos). *** (*p* < 0.001). Animal is to the top in D. Scale bar, 50. (G) Schematic diagram showing the proposed drag-mediated mechanism guiding the motion of attached DFCs. Tension generated by YSL/EVL vegetal spreading is transmitted to DFCs through apical attachments to guide their vegetal movements. Isolated DFCs devoid of apical attachments (detached) are insensible to YSL/EVL tensile forces and show non-directional movements.

### DFCs sustain a collective vegetal movement despite the increase of detached cells

The tracking of single isolated detached DFCs revealed that they failed to engage in directed and persistent vegetal movements (Figures 4D-4G; Supplementary Video 5). Importantly, this behaviour was also present in a subset of detached cells located close to or inside the central cluster of DFCs during epiboly. In about 70% of embryos, we found that one or two detached DFCs left the cell cluster moving mainly in direction to the animal pole being sequestered by the internalisation movement of the DCL, incorporating into this embryonic cell layer (Figures 5A and 5B; Supplementary Video 6). Remarkably, this group of DFCs mimicked the morphology and movement behaviour of endodermal cells and at later stages differentiated into endodermal tissue derivatives (Supplementary Figure 4A). Furthermore, DFCs transplanted into the paraxial region of host embryos, where endodermal and mesodermal progenitors develop, integrated only into endodermal tissue derivatives (Supplementary Figure 4B). Thus, detached DFCs are prone to leave the cell cluster being sequestered by the DCL, and in this new environment integrate into the endodermal cell layer, losing their primal fate. However, this behavior was barely seen in normal development. Indeed, detached DFCs following the endodermal path were absent in 30% of all analysed embryos and they represented only a small fraction of the complete pool of detached DFCs in the remaining 70% of cases (Figure 5A). In contrast, the number of detached DFCs increased steadily during epiboly as cells completed the process of delamination (Figures 3B). These findings prompted us to investigate how the detached DFC population integrates into the collective movement of attached DFCs, which is guided by the extra-embryonic tissues. To address this issue, we first analysed how the detached DFC population evolves during their collective vegetal movement. Immunostaining for ZO-1 and F-actin revealed that at the onset of epiboly the entire population of DFCs were transiting the process of delamination and thus were all attached to the YSL and EVL (Figure 5C). However, as epiboly progressed, detached DFCs steadily increased to outnumber the attached DFC population from 75% epiboly, and by 90% epiboly they became the predominant pool of DFCs (Figure 5C). Through time-lapse microscopy, we found that the progressive increase of detached cells was not only due to DFCs completing the process of delamination (Figure 3B), but detached DFCs also emerged from events of cell division (Figures 5D and 5E). As previously noted, despite the continuous increase in the number of detached DFCs, the entire population of DFCs still moved in direction to the vegetal pole in coordination with the vegetal spreading of extra-embryonic tissues until advanced stages of epiboly (Figures 5E and 5F). Altogether, these results reveal that the DFC cluster retains a collective vegetal motion regardless of the progressive expansion of the detached DFC population, raising the question of how detached DFCs move towards the vegetal pole.

**Figure 5.**
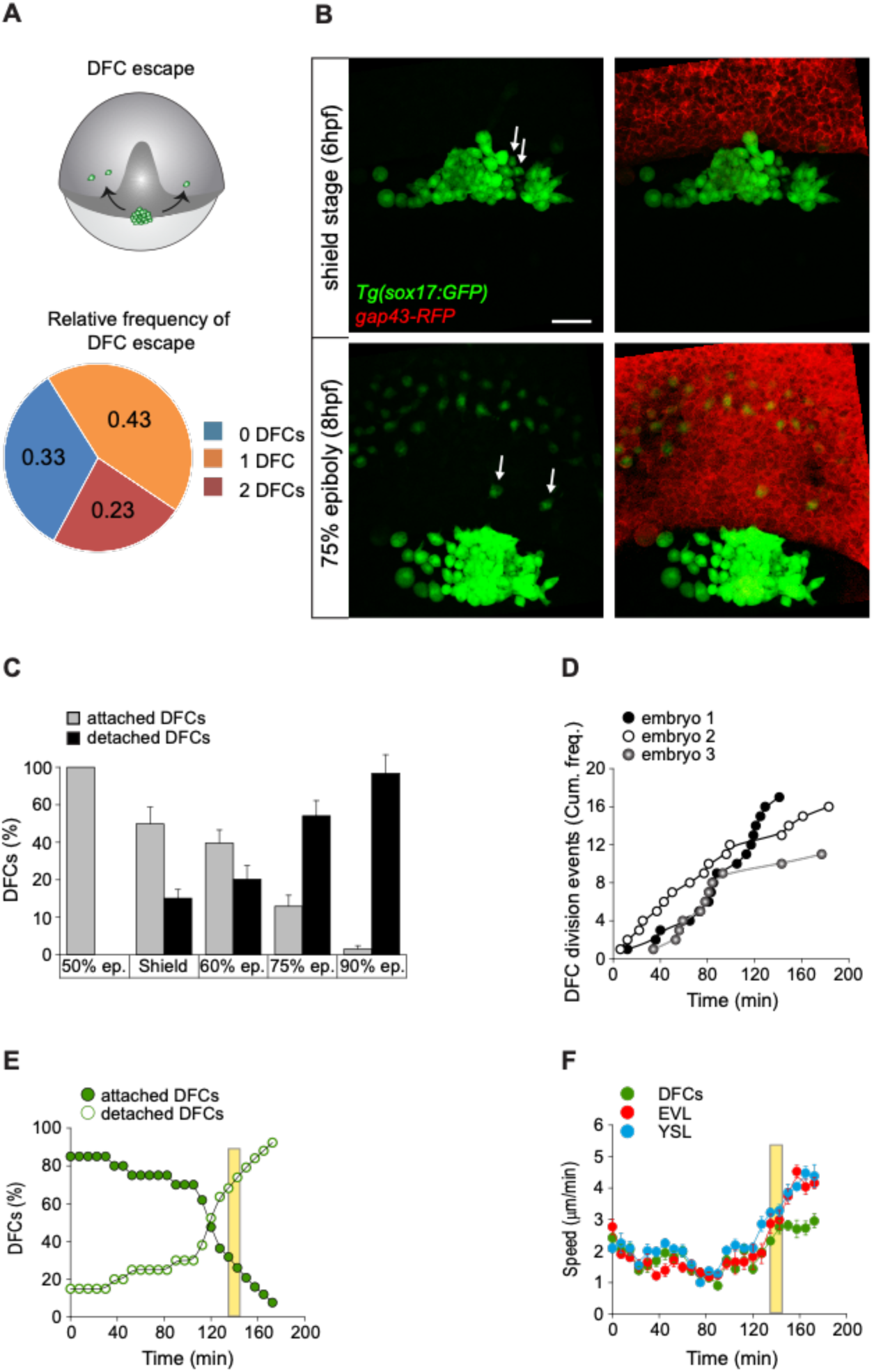
DFCs move to vegetal pole as a collective despite the increase of detached cells. (A and B) Individual detached DFCs can leave the main cluster towards the DCL during normal development. (A) Schematic diagram showing events of escape (top) and quantification of escaped detached DFCs after *in vivo* imaging of Tg(*sox17::GFP*) embryos (bottom, n=30 embryos). (B) Dorsal views of confocal z-stack maximum projections at shield stage (top) and 75% epiboly (bottom) of a representative Tg(*sox17::GFP*) embryo injected with *gap43-RFP* mRNA, labelling DFCs in green and the plasma membrane of all cells in red (extracted from Supplementary Video 6). Arrows indicate two detached DFCs leaving the main cluster towards the DCL. These cells then mimic the behaviour of endodermal cells and differentiate into endodermal tissue derivatives (Fig. S4). Animal is to the top. Scale bar, 50 µm. (C-E) Origin and progressive increase of detached DFCs during development. (C) Quantification of attached and detached DFCs between 50% and 90% epiboly (5.3-9 hpf), as determined from fixed embryos stained with phalloidin and ZO-1 (see Figures 1C and 1D). Values correspond to means ± s.d. (n=10 embryos per stage). (D) Kinetic of cumulative frequency of DFC division events in 3 representative embryos. (E) *In vivo* kinetics of attached and detached DFCs in a representative living Tg(*sox17::GFP*) embryo injected with *lifeactin-mcherry* mRNA to label F-actin. Plot shows the percentage of attached (filled green circles) and detached (empty green circles) DFCs over time. (F) *In vivo* progression in the movement speed of the DFC cluster (green), EVL (red) and the actomyosin ring of the YSL (blue) in the same embryo as in E. In E and F, movies started at shield stage and extended until 90% epiboly. The vertical yellow bar indicates the stage when the movement of DFCs uncouples from the vegetal movement of the EVL and YSL (around 80% epiboly, when detached DFCs reach ∼80%).

### DFC-DFC contact interactions integrate detached cells to the vegetal movement of attached DFCs providing a clustered collective movement

The observation that most detached DFCs move towards the vegetal pole despite their ability to migrate away from the DFC cluster and being sequestered by the DCL, suggests that specific mechanisms integrate these detached cells to the movement of attached DFCs. During epiboly, DFCs transit from being a collection of scattered progenitors into a tight cellular cluster (Figure 1A; Supplementary Video 1) (Oteiza, et al., 2008). DFCs express *e-cadherin* (Figure 6A) (Kane, et al., 2005) and previous reports have shown that functional abrogation of this adhesion molecule affects the cluster cohesion of DFCs leading to a scattered organisation of DFCs at 90% of epiboly (Oteiza, et al., 2010). Thus, DFC-DFC adhesion mediated by E-cadherin (E-cad) could fulfil the role of integrating detached DFCs to the vegetal movement of attached DFCs. Therefore, we tested by time-lapse microscopy if the directed vegetal movement of detached DFCs requires contact interactions mediated by E-cad. We found that E-cad knockdown disrupted cluster cohesion, leading to the creation of multiple small racemes of DFCs that moved in the vegetal direction guided by their apical attachments with the YSL and EVL (Figure 6B). Remarkably, the number of detached cells leaving the DFC collective towards the DCL increased significantly in these embryos (Figures 6B and 6C). These results indicate that DFC-DFC adhesion mediated by E-cad is required for the recruitment of detached DFCs into the vegetal motion of attached DFCs, promoting the formation of a tight cellular cluster.

**Figure 6.**
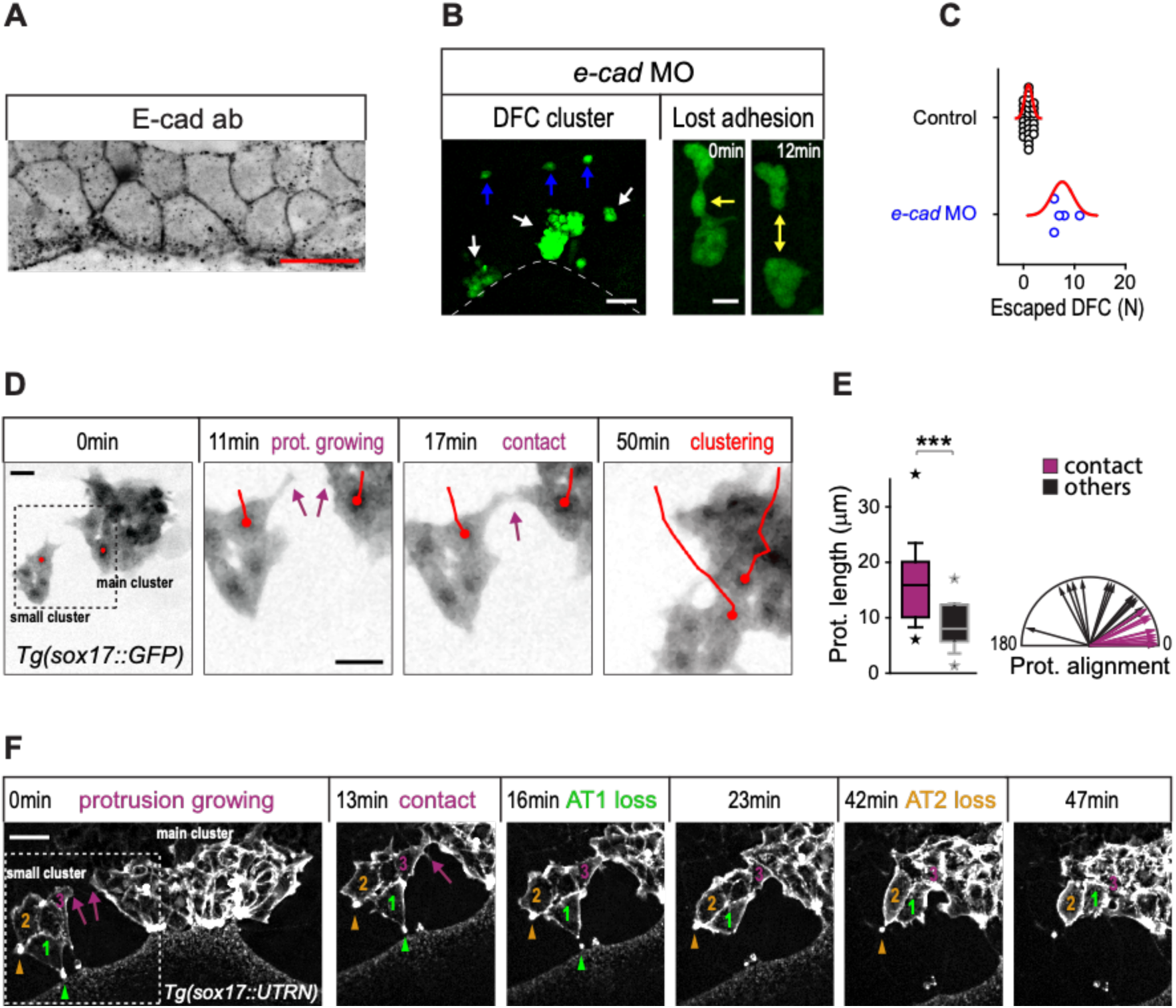
Contact interactions between DFCs couple the motion of attached and detached cell promoting a clustered collective movement. (A) Single confocal plane view of E-cad immunostaining in DFCs in a representative 60% epiboly embryo. Scale bar, 20 µm. (B) Dorsal views of confocal z-stack maximum projections of DFCs from representative Tg(*sox17::GFP*) embryos injected with *e-cad* MO showing the defective collective organisation into multiple racemes (left, white arrows), the increase of escaped cells (left, blue arrows) and the loss of DFC-DFC adhesion contacts (right; arrow indicates the contact and double-arrow the loss of contact). Scale bars, 50 µm (left) and 20 µm (right). (C) Distribution of DFC escape events observed in control and *e-cad* MO injected embryos (n=5 for *e-cad* MO, n=30 for controls). (D) Temporal confocal series of DFCs from a Tg(*sox17::GFP*) embryo showing long protrusions (purple arrows) contacting DFCs from neighbouring clusters. After the initial contact, DFCs from the lateral small cluster approach and establish adhesive contacts with the main cluster (red lines). Scale bars, 20 µm. (E) Quantification of the length (left) and directionality angle (right) of long protrusions involved in DFC-DFC contacts (purple, n=5 embryos) and other protrusions (black, n=5 embryos). *** (*p* < 0.001). (F) Temporal confocal series from a representative Tg(*sox17::utrn-GFP*) embryo showing a DFC from a small lateral cluster (cell 3) sending long protrusions and establishing adhesive contacts with the main cluster before other DFCs from the small cluster (1 and 2) lose apical attachments (arrowheads) (extracted from Supplementary Video 8). Scale bar, 20 µm.

Nonetheless, it remained obscure how cells located at long distances in the initially dispersed group of DFCs approached to each other to establish adhesive contacts. To address this question, we followed the movement of DFCs by *in vivo* microscopy. Our time-lapse image analysis revealed that distant DFCs initiated adhesive contacts by sending long polarised protrusions (Figures 6D, 0 to 17 min, and 6E; Supplementary Video 7). These protrusions resulted in the establishment of long-term adhesion among cells initially separated up to a distance of 52.67±14.72 (mean ± S.D; n=6 embryos), allowing the clustering of DFCs (Figure 6D, 50 min; Supplementary Video 7). Thus, long-range interactions mediated by protrusions work like seeds for the establishment of stable inter-DFC adhesion. Remarkably, using transgenic embryos expressing F-actin only in DFCs we observed a temporal connection between formation of protrusion-mediated adhesive contacts and the loss of apical attachments. Subsets of scattered DFCs that contained several cells with apical attachments started to detach from the extra-embryonic tissues only after initiating protrusion-mediated adhesive contacts with the central cluster of attached DFCs (Figure 6F, 0 to 16 min; Supplementary Video 8). Subsequently, they completed the process of delamination when they became fully integrated into the central DFC cluster (Figure 6F, 23 to 47 min; Supplementary Video 8). Collectively, these findings indicate that cell-cell contact interactions among DFCs integrate detached cells to the vegetal movement of attached cells to guide the clustering and directional vegetal movement of the entire DFC collective.

## Discussion

Here we show a previously unexplored mechanism of morphogenesis of a small group of progenitor cells that stems from the mechanistic link between the process of epithelial delamination underlying progenitor specification and the directed movement of adjacent extra-embryonic tissues. This morphogenetic mechanism safeguards progenitors from undesired losses while guiding their directed motion and allocation as a cluster at the site of organ differentiation. In zebrafish, DFCs are the laterality organ progenitors and arise by delamination from the EVL, an extra-embryonic surface epithelium that protects the early embryo and at later stages gives rise to the periderm (Kimmel, et al., 1990). When DFCs are formed, the EVL epithelium spreads from the equator to the embryo’s vegetal pole during the movement of epiboly in conjunction with the YSL, an extra-embryonic syncytium to which the vegetal margin of the EVL is tied. Remarkably, the apical constriction process underlying DFC delamination follows a temporal progression that allows DFCs to retain long-term apical attachments with the EVL and YSL as they spread towards the vegetal pole during epiboly. Apical attachments work as tissue connectors that couple DFCs with the vegetal spreading of extra-embryonic tissues, guiding their motion towards the site of differentiation at the vegetal pole. In contrast with previous hypotheses indicating that autonomous motility (Ablooglu, et al., 2010) and contact-mediated repulsive interactions with the marginal DCL (Zhang, et al., 2016) drive DFC vegetal movement, we show that mechanical drag by extra-embryonic tissue movement provides the critical guidance cues for DFC directed locomotion. Remarkably, this guidance mechanism is mediated by apical attachments that stem from an incomplete process of delamination, allowing DFCs to coordinate their movement with the adjacent extra-embryonic tissues. Epithelial delamination is a conserved mechanism to generate new mesenchymal cell types in various developmental contexts (Thiery, et al., 2009; Shook and Keller, 2003). Here we show that besides this canonical function, incomplete delamination serves as a generic mechanism for coordinated tissue movement during development, driving the allocation of newly formed mesenchymal cell groups.

Apical attachments guiding DFC movement arise from a process of apical constriction that depends on actomyosin contractibility and is asynchronous among DFCs. We show that such asynchrony generates two distinct populations of progenitors, one holding apical attachments and being pulled by the vegetal movement of extra-embryonic tissues, and a second population of detached delaminated DFCs that follows the vegetal movement of apically-attached DFCs through cell-cell contact mechanisms. In the context of these evolving two populations, having an asynchronous process of cell delamination potentially increases the chances of maintaining a minimal number of apically-attached DFCs able to carry the detached DFC population along their movements, a factor that becomes relevant as epiboly progresses and the ratio of attached/detached DFCs decreases. Furthermore, as apical constriction imposes mechanical stress along the epithelial plane (Heer and Martin, 2017; Martin and Goldstein, 2014) we can speculate that having asynchrony in a collective process of apical constriction promotes the even dissipation of mechanical stress over time and space, protecting the integrity of the epithelium. Future work combining mechanical perturbations/measurements with physical modelling will have to test these hypotheses directly.

Delaminated DFCs are intrinsically motile and can move towards the DCL, being sequestered by the massive internalisation movements of this embryonic cellular domain during gastrulation. Importantly, when DFCs reach the DCL, either during normal development or after transplantation, they follow the endodermal path revealing that DFCs have a previously unrecognised potential to become endoderm that is expressed if they enter the developmental field of the endoderm. Such potential co-option of DFCs by the endodermal DCL reduces the number of progenitors, and this can have a detrimental impact on left-right asymmetry development, increasing the incidence of embryo laterality defects (Moreno-Ayala, et al., 2020). Here we show that DFCs transiting the process of delamination avoid endodermal fate by establishing E-cadherin mediated adhesive contacts with the sub-population of attached DFCs before completing the process of delamination. Thus, DFC-DFC contact interactions mediated by E-cadherin play a dual function, to protect delaminated cells from escaping towards the endoderm and to ensure they move together with the attached DFC population as a collective group. Importantly, DFCs located at long distances establish adhesive contacts by sending long polarised protrusions whose persistence and directionality differ from the short random protrusions formed by most DFCs at the edge of the DFC cluster. The mechanisms underlying the formation of these directed long protrusions are currently unknown. Notably, recent work shows that migrasomes containing the chemokine ligands Cxcl12a/b become enriched in the extracellular space surrounding DFCs (Jiang, et al., 2019) thus opening the possibility that long polarised protrusions are a manifestation of a chemoattraction mechanism. It is also remarkable the observation that sub-clusters of DFCs delaminate only after establishing adhesive contacts with the main DFC cluster suggesting a mechanistic coupling between the cell adhesion, cell motility and the process of delamination, which will be interesting to explore in future work. Together, these findings provide a novel developmental function for E-cadherin, mediating the transfer of movement information between two distinct populations of progenitor cells defined by their state in the delamination process: attached (transiting delamination) and detached (delaminated). Remarkably, motion transfer from attached to detached DFCs resembles the leader-to-follower motion transmission observed in events of collective cell locomotion, many of which also require cadherin-dependent cell-cell contact interactions (Reig, et al., 2014; Theveneau and Mayor, 2013; Friedl and Gilmour, 2009). In the case of DFCs, the role of E-cadherin in motion transmission complements the previously described function in clusterisation (Matsui, et al., 2011; Oteiza, et al., 2010; Hong and Dawid, 2009), and together ensure that progenitors reach the site of terminal differentiation in a sufficient number and organised as a tight collective, a prerequisite to proceed into further stages of organogenesis (Oteiza, et al., 2010; Oteiza, et al., 2008).

Developmental cases of cells being dragged through physical bonds with adjacent tissues have recently begun to be reported. Among them, *C elegans* primordial germ cells internalise during gastrulation due to cohesive contacts that these cells establish with the moving ingressing endoderm (Chihara and Nance, 2012). In annual killifish, the embryonic DCL spreads during epiboly as a result of adhesive contacts with the basal epithelial domain of the expanding EVL (Reig, et al., 2017). In zebrafish, the vegetal spreading of the EVL during epiboly follows the autonomous movement of the YSL to which is tightly bonded at its margin by TJ complexes (Schwayer, et al., 2019; Behrndt, et al., 2012; Koppen, et al., 2006; Betchaku and Trinkaus, 1978). Therefore, mechanical drag is an emergent cell movement mechanism whose extent and impact during embryo morphogenesis needs to be further investigated. Here we show that drag-mediated locomotion is a crucial driver of organogenesis that emerges at the interface of embryonic and extra-embryonic cellular domains highlighting the essential role of mechanical information from extra-embryonic tissues in driving early embryo development (Christodoulou, et al., 2019; Reig, et al., 2017; Hiramatsu, et al., 2013).

## Acknowledgements

We thank the bioimaging and zebrafish facilities of ICBM-U.Chile and IST Austria for continuous support. We also thank Felipe Santibañez and Mauricio Cerda for providing algorithms for image analysis.

## Material and Methods

### Fish Strains and Maintenance

Zebrafish (*Danio rerio*) strains were maintained and raised according to previously published procedures (Westerfield, 2000). Embryos were grown in E3 solution at 28°C and staged according to morphology. Fish care and procedures were approved by the Ethical Review Committee and comply with the Animals Scientific Procedures Act 0466. Zebrafish strains used were: wild type AB, Tg(*actb1::myl12.1-eGFP*) (Behrndt, et al., 2012), Tg(*sox17::utrn-GFP*) (Woo, et al., 2012), Tg(*sox17::GFP*) (Sakaguchi, et al., 2006), Tg(*actb1::mCherry-utrCH*) (Behrndt, et al., 2012) and Tg(*actb1::myl12.1-eGFP; actb1::mCherry-utrCH*).

### Morpholino and mRNA injections

Synthetic mRNA was produced using the SP6 mMessage mMachine kit (Thermo Fisher Scientific). Glass capillaries (BF100-98-15, Sutter Instruments) were pulled using a needle puller (P-97, Sutter Instruments) and mounted on a microinjection system (Picospritzer III, Parker Hannifin). Embryos were microinjected at the 1-cell stage as previously described (Barth and Wilson, 1995), unless stated otherwise. 100 pg of *Gap43-RFP* (Reig, et al., 2017) or 40 pg of *lifeACT-RFP* (Behrndt, et al., 2012) mRNA or *h2b-GFP* (Keller, et al., 2008) were injected as a counterstain for whole embryo visualisation. 20 pg of *zo1-GFP* mRNA was injected to label apical junctions during DFC delamination (Schwayer, et al., 2019). 75 pg of *N-ter(1-300aa)-Mypt1* mRNA (Jayashankar, et al., 2013) was injected into the yolk cell at 3.3 hpf for functional inhibition of the actomyosin network in the YSL. 2 ng of *cdh1* MO (5’-TAAATCGCAGCTCTTCCTTCCAACG -3’, GeneTools) (Maitre, et al., 2012) was injected to abrogate *cdh1* function.

### Immunohistochemistry

Embryos between 50% and 90% of epiboly (5.3-9 hpf) were fixed and stained as described previously (Oteiza, et al., 2008). Embryos were mounted on agarose-coated dishes embedded in 1% low melting-point agarose. Samples were imaged on a Leica TCS LSI Confocal microscope with HCS software using a 5x objective and 488/520 (λexc/λem) lasers. The following antibodies and dilutions were used: mouse anti-ZO-1 (339100 Invitrogen, 1:200), anti-pMLC2 (3671 Cell Signaling, 1:200), anti-Cdh1 (MPI-CBG #174, 1:200), goat anti-mouse Alexa Fluor 488 (A-11001, Thermo Fisher Scientific) and goat anti-rabbit Alexa Fluor 568 (A-110011, Thermo Fisher Scientific).

### Whole embryo confocal imaging

Tg(*sox17::GFP*) embryos injected with 50 pg of *gap43-RFP* mRNA were mounted in 0.5% low melting point agarose in a custom designed chamber at either dome or 50% epiboly stage. The temperature was kept constant at 28°C throughout the imaging experiment using a temperature control system. Whole embryo *in vivo* microscopic imaging was performed in a Leica TCS LSI Confocal microscope with HCS software using a 5x objective and 488/520 (λexc/λem) lasers.

### High-resolution confocal imaging

Tg(*sox17::GFP*) and Tg(*sox17::utrn-GFP*) embryos were used for high-resolution confocal imaging of cell protrusions. Tg(*actb1::myl12.1-eGFP*) and *lifeactin-RFP* injected embryos were used to analyze myosin II and F-actin *in vivo*. Embryos were imaged from shield stage onwards in a Volocity ViewVox® spinning disc (Perkin Elmer®) coupled to a Zeiss Axiovert 200 confocal microscope using a Plan-Apochromat 40x/1.2W lasers 488/520, 568/600 and 647/697 nm (λexc/λem). Processing and analysis of digital images were performed using Fiji (Schindelin, Arganda-Carreras et al. 2012), Matlab (Matlab 2014), Volocity (Improvision®) and Adobe photoshop.

### Laser ablation

Mechanical disruption of the actomyosin ring within the YSL was performed as previously described by conducting laser ablation on a UV laser ablation setup (Behrndt, et al., 2012) equipped with a Zeiss 63x 1.2 NA water immersion lens using Tg(*actb1::myl12.1-eGFP*) embryos. Embryos were mounted at 50% epiboly (5.3 hpf) and the YSL actomyosin cortex close to the EVL margin was repeatedly ablated by applying 10 UV pulses at 1,000 Hz on a rectangular ROI. Suboptimal ablation intensity was applied to just disrupt the cortical actomyosin flux and marginal ring, and avoid the activation of a wound response within the yolk cell. The kinetic of EVL cells, DFCs and YSL marginal actomyosin network adjacent to the disrupted cortex was compared with the kinetic of close neighbouring tissues showing intact regions of the actomyosin network as an internal control. Cortical laser ablation of the EVL was performed parallel to the EVL margin and perpendicular to the EVL actomyosin cortex by applying 25 ultraviolet pulses at 1,000 Hz along a 10μm-line. Retraction of apical ties and DFCs were quantified from maximum *z*-projections images.

### Actomyosin dynamics analysis

Images obtained from *in vivo* imaging of Tg(*actb1::myl12.1-eGFP*) embryos were segmented manually as described below in the method section to obtain the apical ROI of DFCs. A maximum z-projection of 5 μm of depth using Fiji was applied to each xyz stack to obtain a 2D sequence. Florescence intensity was calculated in Fiji as the average intensity per pixel in each apical ROI during the entire process of apical constriction underlying DFC delamination.

### Cell protrusion analysis

Quantification of cell protrusions was assessed by making volumetric binary masks from 4D confocal images using manual threshold to keep the presence of protrusions and avoid artefacts. Each volume was reduced to a 2D cluster mask by z-projection based on maximum intensity. Manual correction allowed to keep protrusion zones that were occluded due to dimensional reduction. The Fiji tool Local Thickness was used to generate a central mask to describe the cluster central zone without the protrusions. Masks containing just the protrusions zones were obtained by subtracting cluster masks and central masks. Each protrusion was defined with a protrusion axis that connected the base and tip of the protrusion. A long axis connected the cluster centroid and the base of each protrusion. Individual protrusion orientation was described calculating the angle between the long axis and the protrusion axis.

### Cell segmentation

Images obtained from *in vivo* imaging of *gap43-RFP* or *ZO1-GFP* mRNA injected embryos were segmented manually using Wacom Cintiq Touch Screen tablet (Wacom) and Fiji software (Schindelin, et al., 2012). A maximum z-projection using Fiji was first applied to each xyz stack to obtain a 2D sequence and simplify the xy segmentation of cell boundaries. To perform segmentation in the yz plane, z-projections containing the central region of cell volume of marginal and submarginal DFCs were selected and reconstructed using Fiji.

### Cell tracking

Cell movement was tracked by following the cell’s center in Tg(*sox17::GFP*) embryos, and the nucleus in *H2B* mRNA injected embryos. YSL tracking was performed by following arbitrary landmarks inside the marginal actomyosin network. Tracking of apical attachments was assessed by following actin rich zones localised in the apical face of DFCs. Tracking was performed manually in 2D from maximum z-projections using Fiji plugin MTrack J. Speed and persistence (ratio of displacement to trajectory length) of movement was then calculated from tracking data.

### Statistical analysis

All experiments were performed at least three times. Unless indicated, plots display the mean and the standard deviation. Statistical inference analysis was conducted initially by a *Shapiro-Wilk* test to assess if the data were normally distributed. Two-sample *F*-test of equality of variance was applied to assess if the samples had the same variance. Significance for two groups with normal distribution was calculated through a two-tail *t*-test and the *p*-value was selected depending if the dataset had or not equality of variance. For other distributions, a non-parametric *Kolmogorov-Smirnov* test was applied. A one-way ANOVA test was conducted to compare more than two groups. If the ANOVA test indicated significant differences between the means a *Bonferroni* test was conducted to calculate the level of significance between the samples. All statistical analyses were conducted using the Origin 2016 (OriginLab).

## Key Resource Table

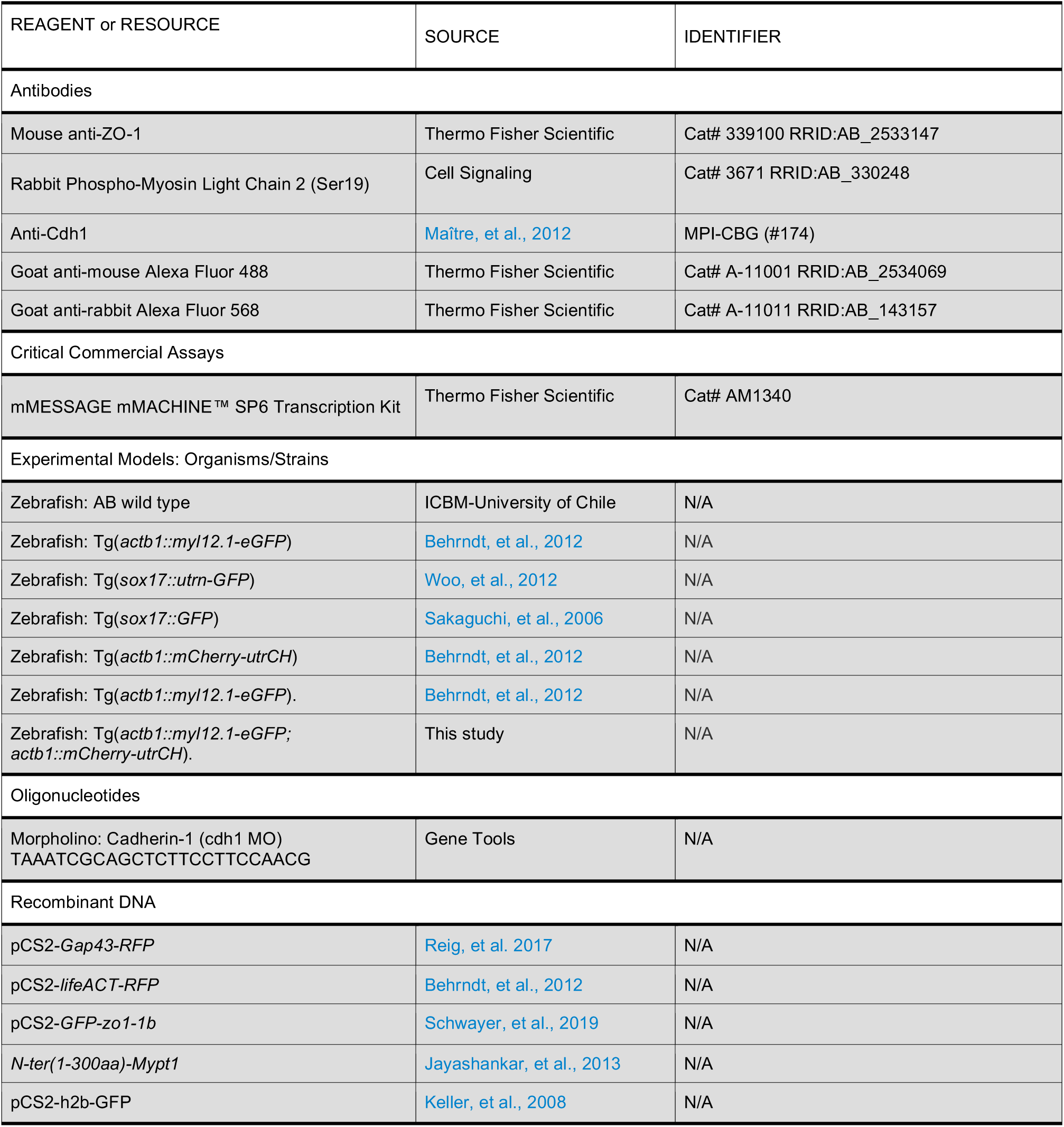

| Software and Algorithms |  |  |
| --- | --- | --- |
| Fiji | Schindelin, Arganda-Carreras et al. 2012 | https://imagej.net/Fiji |
| MATLAB | MATLAB Software | https://la.mathworks.com/products/matlab.html |
| Volocity | Quorum Technologies Inc | https://quorumtechnologies.com/ |
| Origin | OriginLab | https://www.originlab.com/ |

## Supplemental Information

### Supplemental Figures

**Supplementary Figure 1.**
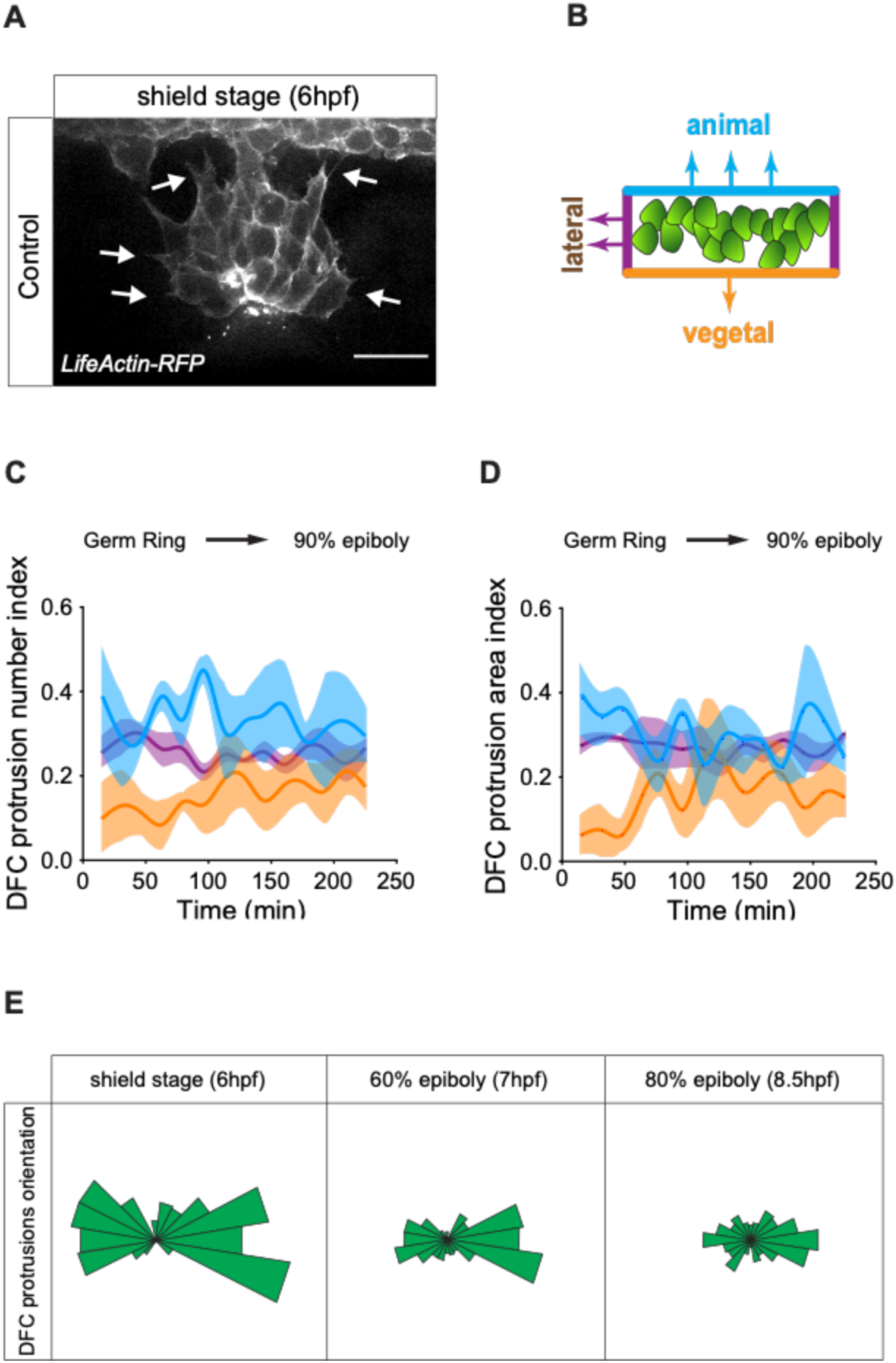
DFC protrusions are not polarised in direction of the vegetal movement of the cluster. (A) Dorsal views of DFCs from a living *lifeactin-RFP* injected embryo at shield stage, with animal to the top. White arrows indicate lamellar- and filopodial-like cell protrusions. Scale bar 50 µm. (B) Scheme of the DFC cluster showing how cell protrusions extending from the vegetal (orange), lateral (purple) and animal (light blue) edges of the cluster were quantified in Tg(*actb1:lifeactin-RF*P) embryos to build the plots of C and D. (C) Kinetic of DFC protrusion number index (n=3 embryos). (D) Kinetic of DFC protrusion area index (n=3 embryos). (E) Circular distribution plots of DFC protrusions at different developmental stages obtained from fixed Tg(*sox17:utrn-GFP*) embryos (n=10 per developmental stage).

**Supplementary Figure 2.**
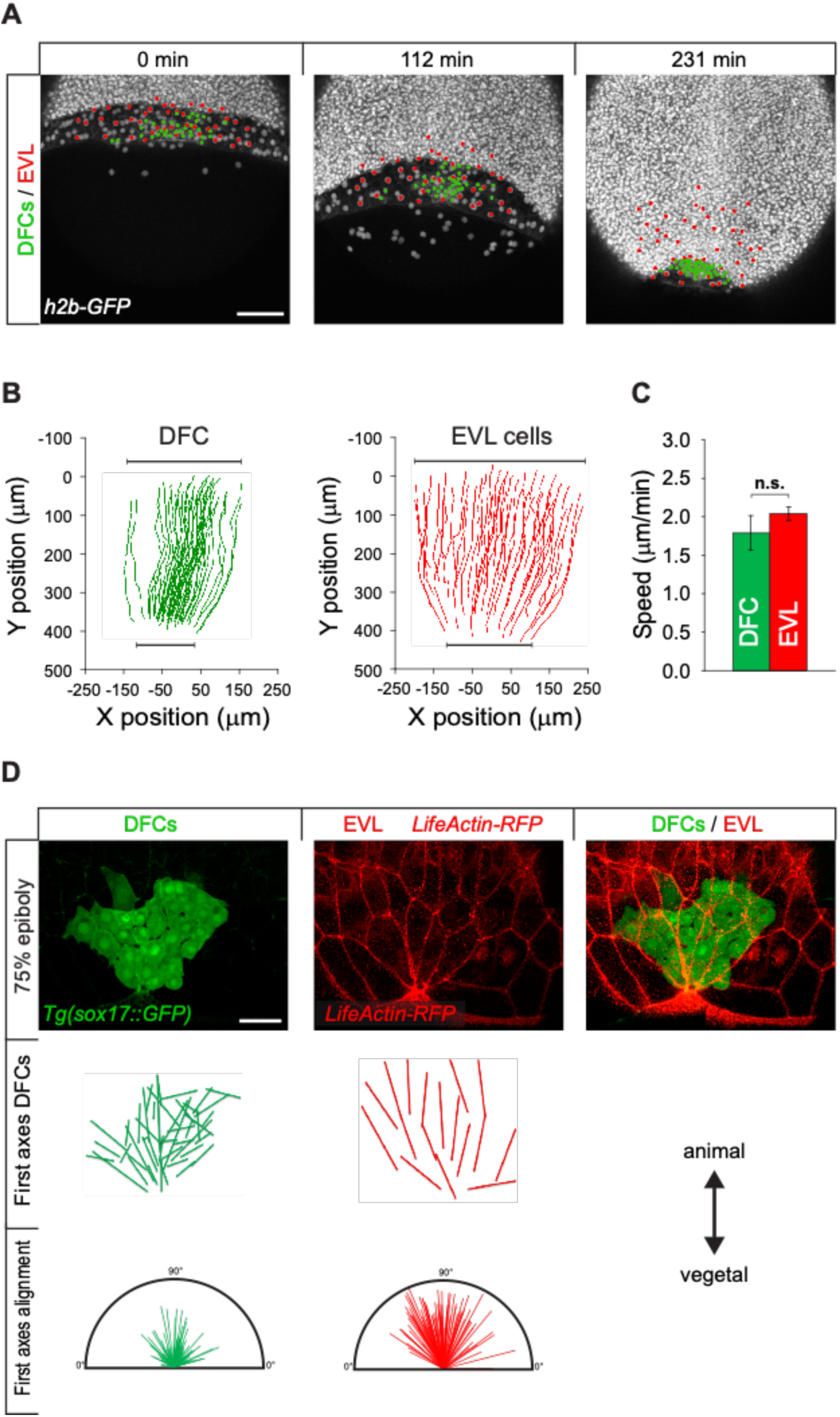
The movement of DFCs mirrors the pattern of EVL vegetal spreading. (A) Dorsal views of confocal z-stack maximum projections of a wild type embryo injected with h2b-GFP mRNA to label all nuclei (white dots in left and middle panels), and the corresponding positions of nuclei showing the concordant movement of DFCs (green dots) and overlying dorsal EVL cells (red dots). Dorsal view with animal to the top, from shield stage (6 hpf). Scale bar, 100 µm. (B) Cell tracking of the vegetal movement and convergence between shield and 90% of epiboly of DFCs (green) and overlying EVL (red) in a representative wild type embryo injected with *h2b-GFP* mRNA. (C) Speed of DFCs (green) and EVL (red), expressed as means ± SD (n = 3 embryos). n.s. (non significant). (D) Dorsal views of DFCs, EVL cells and merge from a representative living Tg(*sox17:GF)* injected with *lifeactin-RFP* embryo at 75% of epiboly (top). Spatial representation of the alignment of the first principal axis of DFCs and EVL cells (middle). Distribution plot of the alignment of first principal axis of DFCs and EVL cells from 3 embryos (bottom). Animal is to the top in all panels. Scale bar, 50 µm.

**Supplementary Figure 3.**
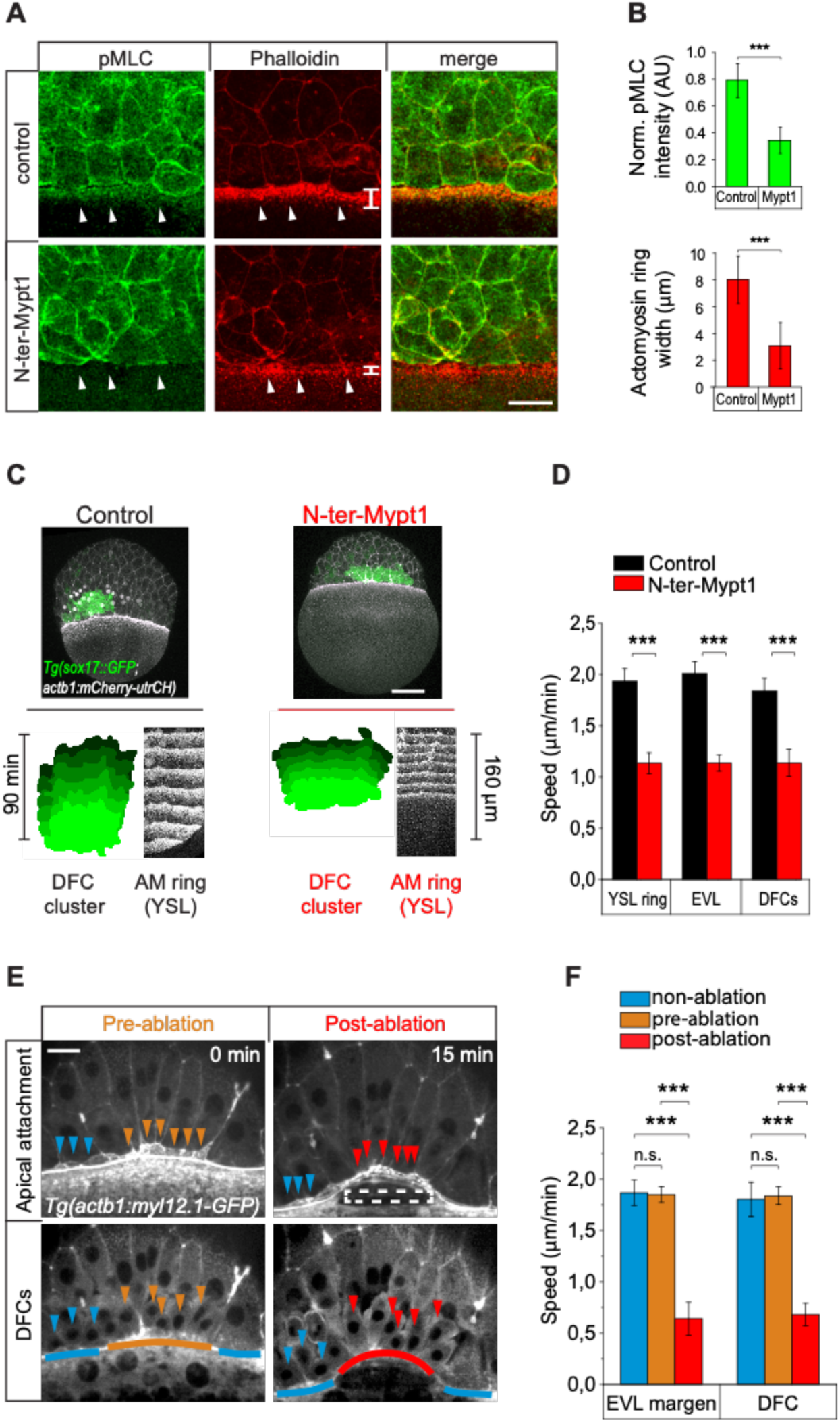
Disruption of the YSL actomyosin network impairs DFC vegetal movement. (A and B) The YSL actomyosin ring is disrupted in embryos over expressing N-ter-Mypt1 in the yolk cell. (A) Immunostaining of phospho-myosin light chain II (left), phalloidin (middle) and merge (right) at 75% of epiboly in controls (top) and embryos injected with N-ter-Mypt1 specifically in the yolk cell (bottom). White arrows indicate the actomyosin ring in the YSL that is affected in the injected embryos. Animal is to the top. Scale bar, 20 μm. (B) Quantification of phospho-myosin light chain II fluorescence intensity (top) and actomyosin ring width (bottom) in control and N-ter-Mypt1 injected embryos. Values correspond to means ± S.D. (n=3 embryos). *** (p < 0.001). (C and D) DFC movements are affected after disruption of the YSL actomyosin ring. (C) Dorsal views of confocal z-stack maximum projections of Tg(*sox17::GFP; actb1::mCherry-utrCH*) embryos at 60% epiboly (7 hpf). Top images correspond to control (left) and experimental embryo injected with 5 pg of *N-ter-Mypt1* mRNA into the yolk cell at 3.3 hpf (right). Bottom kymographs show the movement of DFCs (green) and the YSL actomyosin ring (white) between 60% and 80% epiboly (7-8 hpf) in control (left) and N-ter-Mypt1 (right) conditions. Animal is to the top in all panels. Scale bar, 100 μm. (D) Quantification of the vegetal speed of DFCs, EVL and the YSL actomyosin ring from 60% to 80% epiboly in control (black) and N-ter-Mypt1 (red) conditions. Values correspond to means ± S.D. (n=3 embryos). *** (p < 0.001). (E and F) Laser disruption of the the YSL actomyosin network impairs DFC vegetal movements. (E) Dorsal views of a Tg(*actb1::myl12.1-GFP*) embryo at early shield stage (5.8 hpf) before and after laser ablation of the cortical actomyosin ring of the YSL. Confocal optical planes are at the level of the EVL showing the apical contacts (top) and DFCs (bottom). The dashed rectangle depicts the zone of laser ablation, arrowheads point to apical contacts (top) and individual DFCs (bottom), while lines depict the position of the EVL margin immediately above the ablation zone (orange and red) and lateral to this region (blue). Animal is to the top. Scale bar, 20 μm. (F) Quantification of the vegetal speed of DFCs and the EVL margin in control un-ablated (blue) and ablated zones, prior (orange) and after (red) the ablation. Values correspond to means ± S.D. (n=3 embryos). *** (p < 0.001). n.s. (non-significant).

**Supplementary Figure 4.**
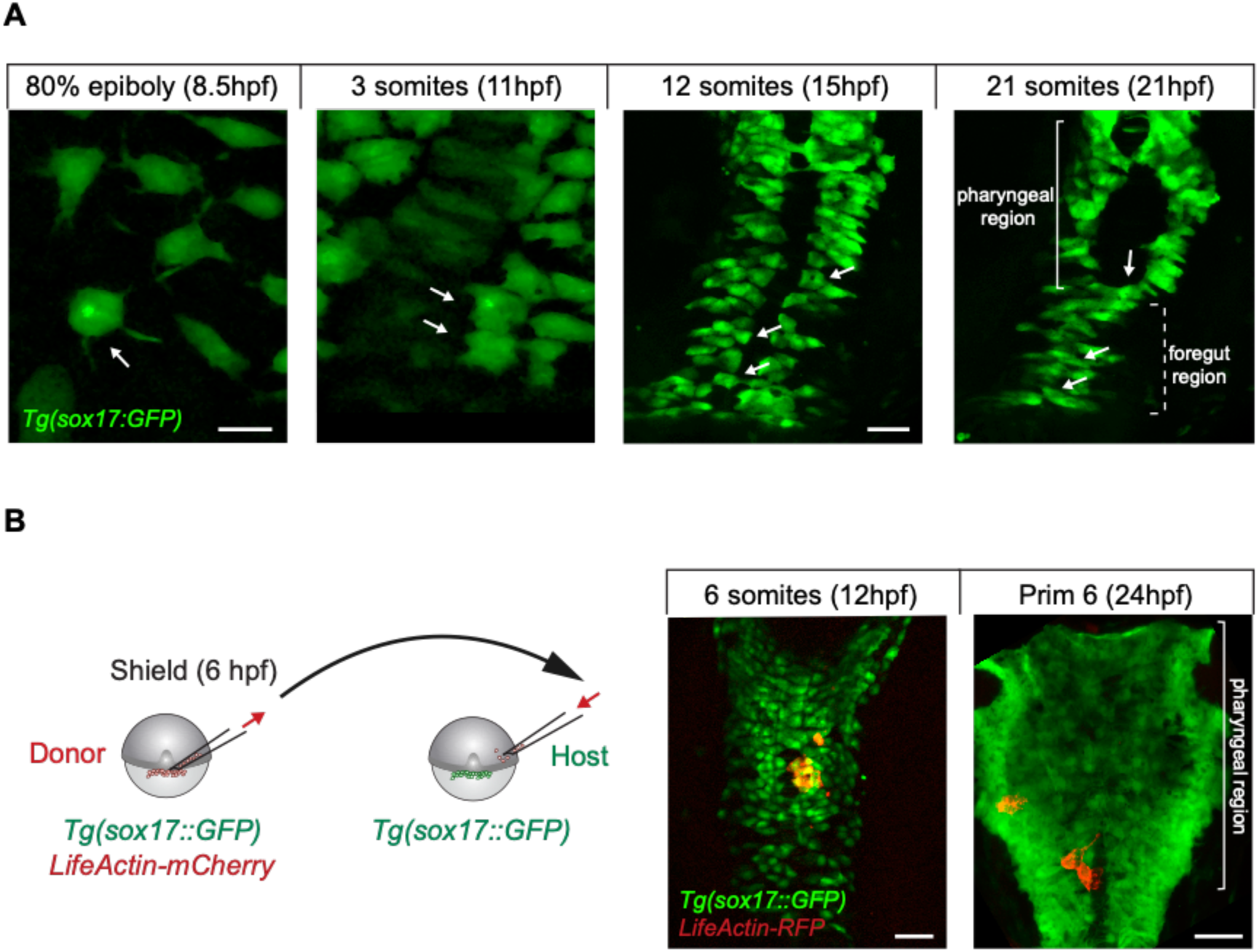
DFCs have endodermal potential and acquire endodermal fate after internalising into the DCL. (A) During normal development, delaminated DFCs can leave the main cluster and internalise into the DCL (see Fig 5A,B). Once in the DCL, these cells mimic the morphology and migratory behaviour of endodermal cells during gastrulation (80% epiboly) and neurulation (3-somites) and later integrate into endodermal tissue derivatives (12- and 21-somites). An example of the behaviour and fate of a escape DFC is indicated with arrows. Scale bar, 20 µm. (B) Isochronic DFC transplantation. DFCs from donor Tg(*sox17:GFP*) embryos injected with *lifeactin-RFP* mRNA were transplanted into the paraxial region of host Tg(*sox17:GFP*) embryos (left). Transplanted DFCs (in red) integrated into endodermal tissue derivatives (in green) (right). Scale bar, 20 µm.

### Legends of Supplementary Videos

**Supplementary Video 1**. **Movement of DFCs from the embryo equator to the vegetal pole during epiboly** (related to Figure 1A). Time-lapse video of confocal z-stack maximum projections of a Tg(*sox17:GFP*) embryo injected with *gap43-RFP* mRNA, showing cytoplasmic GFP (green) in DFCs from early stages (main cell cluster) and later in the forming endoderm (scattered cells), and membrane-tagged RFP (red) in all cells. Dorsal view with animal to the top. The video starts at 50% epiboly (5.3 hpf). Images were acquired every 8 minutes. Scale bar, 100 µm.

**Supplementary Video 2**. **Delamination, apical constriction and vegetal movement of DFCs during epiboly** (related to Figures 1E and 1F). Time-lapse video of confocal z-stack maximum projections of an embryo injected with *zo1-GFP* and *gap43-RFP,* showing dorsal views at the level of the EVL (left, *zo1-GFP* channel) and under the EVL (middle, *gap43-RFP* channel), and the merge image on the right. The surface view shows *zo1-GFP* enriched at the apical junctions of DFCs as they undergo apical constriction. *zo1-GFP* also labels the junctions of dorsal EVL cells with less intensity. DFCs are labelled at time=0 with orange dots. Dorsal EVL cells are labelled at time=0 with blue dots. The middle row is focused under the apical face of EVL and shows *gap43-RFP* membrane staining in delaminating DFCs (orange labelling at time=0), some marginal EVL cells (blue labelling at time=0) and the deep cells of the blastoderm margin (unlabelled at time=0). The bracket indicates the group of ingressing DFCs. The video starts at 50% epiboly (5.3 hpf). Images were acquired every 1.7 minutes. Scale bar, 20µm

**Supplementary Video 3**. **Dynamics of apical constriction of DFCs during spontaneous apical myosin loss and recovery** (related to Figures 2G and 2H). Time-lapse video of confocal microscopy z-stack maximum projections of a Tg(*actb1:myl12.1-GFP*) embryo revealing that apical constriction and relaxation correlates with apical myosin accumulation and loss, respectively. Red arrow indicates the time of maximum apical area relaxation and minimum apical myosin accumulation. Scale bar, 10 µm.

**Supplementary Video 4**. **Apical attachments of DFCs are under pulling tension from extra-embryonic tissues** (related to Figures 4A-4C). Time-lapse video of confocal microscopy z-sections of a Tg(*actb1:myl12.1-GFP*) embryo at 70% of epiboly, focused at the level of the EVL (left panel) and DFCs (right panel), showing the animal-ward recoiling of apical attachments of two DFCs after the laser line ablation of an EVL cortical junction (red line). Dorsal view with animal to the top. Time 0 corresponds to laser ablation, with negative and positive times indicating pre- and post-laser ablation times, respectively. Scale bar, 20 µm.

**Supplementary Video 5**. **Apical attachments promote a persistent vegetal movement of attached DFCs** (related to Figures 4D-4E). Time-lapse video of confocal microscopy z-stack maximum projections of a Tg(*sox17:GFP; actb1:mCherry-utrCH*) embryo expressing cytoplasmic GFP (green) in DFCs and F-actin (white) in all cells. Tracks of isolated DFCs (left panel) reveal that attached DFCs show persistent vegetal movements (light blue cell, and yellow cell before the loss of apical attachment at 50 min). In contrast, detached DFCs devoid of apical attachments (detached DFCs) move with little persistence and lack directionality (red cell, and yellow cell after the loss of apical attachment at 50 min). Dorsal view with animal to the top. The video starts at 60% epiboly (7 hpf). Scale bar, 50µm.

**Supplementary Video 6**. **DFCs can leave the main cluster and internalise into the DCL** (related to Figures 5A and 5B). Time-lapse video of confocal microscopy z-stack maximum projections of a Tg(*sox17:GFP*) embryo (left and middle panels), and the corresponding merge image with bright field (right panel), showing the escape and internalisation into the DCL of two DFCs (light blue tracks). Dorsal view with animal to the top. The video starts at the germ ring stage (5.7 hpf). Images were acquired every 3 minutes. Scale bar, 50 µm.

**Supplementary Video 7. Long polarised protrusions promote the initiation of adhesive contacts between DFCs**. Time-lapse video of confocal microscopy z-stack maximum projections of a Tg(*sox17::GFP*) embryo, with an inverted lookup table, showing DFCs at the edge of the cluster producing long polarised lamellar- and filopodial-like cell protrusions (arrows) that contact distant DFCs and bring them closer to the cluster. Dorsal view with animal to the top. The video starts at 60% epiboly (7 hpf). Scale bar, 20 µm.

**Supplementary Video 8. Polarised protrusions promote the establishment of adhesive contacts and cluster formation before laterality organ progenitors loose their apical** attachments. Time-lapse movie of confocal microscopy z-stack maximum projections of a Tg(*sox17::utrn-GFP*) embryo, showing a small cluster of DFCs in the vicinity of the main DFC cluster. DFCs at the edge of both clusters send long polarised lamellar- and filopodial-like cell protrusions (arrows) and establish adhesive contacts. DFCs from the small cluster then fuse within the main cluster before loosing their apical ties (arrowheads). Dorsal view with animal to the top. The video starts at 70% epiboly. Scale bar, 20 µm.

